# Luteal progesterone level correlated with immunotherapy success of patients with repeated implantation failures

**DOI:** 10.1101/603720

**Authors:** Omar Sefrioui, Aicha Madkour, Nouzha Bouamoud, Ismail Kaarouch, Brahim Saadani, Saaïd Amzazi, Henri Copin, Moncef Benkhalifa, Noureddine Louanjli, with the Lorem Ipsum Consortium

## Abstract

Immunotherapy using PBMC administration demonstrated relatively its effectiveness to treat RIF patients but it still unclear to explain some miscarriages. Luteal progesterone level (LPL) issued from corpus luteum after embryo implantation stage could be informative basis data to personalize immunotherapy for RIF patients predicting clinical outcomes. This randomized controlled study included 70 patients undergoing ICSI program presenting at least 3 RIF: 39 for Control of untreated patients and 31 for PBMC-test concerning treated patients with immunotherapy. For PBMC-test group, Peripheral Blood Mononuclear Cells (PBMCs) were isolated from patients on ovulation induction day and cultured three days to be administered to intrauterine cavity of patients two days before fresh embryo transfer. LPL was analyzed at day 15 after embryo transfer and clinical outcomes were calculated including implantation, clinical pregnancy and miscarriage rates. Clinical outcomes were doubly improved after immunotherapy including implantation and clinical pregnancy rates comparing Control versus PBMC-test (10% and 21% vs 24% and 45%). In the other hand, this strategy showed an increase over double in LPL (4ng/ml for Control vs 9ng/ml for PBMC-test) while the latter was correlated to clinical pregnancy. Bypassing the effectiveness of this immunotherapy approach for RIF patients, it is directly correlated to LPL proving the interactive reaction between immune profile of the treated patients and progesterone synthesis by corpus luteum.

## Introduction

Though in vitro fertilization (IVF) success is generally limited to 30% depending on embryo implantation, the major part of implantation establishment is bypassing embryo quality and its genetic integrity highlighting the communication between embryo and the mother. This cross talk is essentially orchestrated by hormonal and immune dialogues in order to assure embryo invasion in the maternal endometrium without rejecting the fetal allograft. Furthermore, it is already known that progesterone (P4) is one of the most important implantation/pregnancy success keys for its effects on the endometrium and early pregnancy survival while its removal results in miscarriage [1–3].

P4 is a hormonal key to modulate the maternal immune system by reducing natural killer (NK)-cell activity [4], inhibiting cytotoxic T-cell activity [5], increasing HLA-G production in trophoblast cells [6], increasing suppressor-cell levels [7] and as special mechanism, it is able to induce lymphocyte-blocking proteins production such as progesterone-induced blocking factor (PIBF) [8–11]. Generally, its anti-inflammatory effect reported by several studies showing that it is essential to modify the cytokine response from pro-inflammatory profile presented by Th1 during embryo implantation to anti-inflammatory profile presented by Th2 for pregnancy maintain [12–16, 18, 19]. Thus, Th1/Th2 unbalance could explain implantation failures in some patients with RIF, RPL or recurrent miscarriages (RM) [20–22].

Indeed, with special interest on immunology of reproduction, Yoshioka et al. [23] was the first team which could to realize immunotherapy for patients with repeated IVF failures based on intrauterine administration of peripheral blood mononuclear cells (PBMC). Then, several studies developed this novel approach with some modifications on PBMC preparation protocol, including fresh or frozen cycles, embryo day 3 or blastocyst transfer, patients with at least 2, 3 or 4 RIF in order to improve significantly clinical outcomes and decrease the miscarriage rates [22, 24, 25].

Furthermore, PBMC immunotherapy could to be efficient for some RPL cases and avoiding miscarriages [22, 26, 27] despite lack of clear definition toward RIF and RPL [28]. This issue led us to wonder about mechanism of PBMC immunotherapy by what it could modulate maternal immune response homing Th1/Th2 balance required for implantation and pregnancy stage in order to manage RIF and RPL clinical cases and its effect on luteal P4 synthesis. Indeed, it is known that P4 is an indispensable factor for endometrium decidualization and for early stage of clinical pregnancy to prepare an adequate immune environment for fetus and low LPL results abnormal ongoing pregnancy [18, 19, 29–32].

Forward, it is increasingly clear that embryo implantation is dependent in one side on immune local mechanisms and in another side on endocrine mechanisms related to luteal P4 synthesis while their interactive reaction is still kept into question for pregnancy achievement. For this reason, while our previous work [22] was based on the implementation of PBMC immunotherapy for RIF patients and proving its efficiency tripling the clinical outcomes, the present work was more focused on demonstrating the correlation between LPL and immunotherapy success highlighting the interactive reaction between luteal P4 synthesis by CL and immune system.

## Materials and methods

### Ethical Standards

The study was approved by the ethics committee, (Comité d’Ethique pour la Recherche Biomédicale- Faculty of Medecine and Pharmacy, University Mohammed V, Rabat, Morocco) and patients provided written informed consent after being presented with the terms and issues of the study. The authors assert that all procedures contributing to this work comply with the ethical standards of the relevant national and institutional committees on human experimentation and with the Helsinki Declaration of 1975, as revised in 2008.

### Patient’s selection and study design

This was a prospective randomized study over two years conducted in Anfa Fertility Center including 70 couples who attended IVF program with at least 3 RIF without female age limit while 48 patients of them were less than 40 years old. In the selected couples, women had unremarkable clinical history and comparable clinical features and embryological outcomes (Table 1). All women received the same antagonist ovarian stimulation protocol [22] to minimize the effect of other parameters. Indeed, the whole lot was divided into two groups; treated group with PBMC immunotherapy (PBMC-test, n=31) and the control group without treatment (Control, n=39).

**Table 1.**
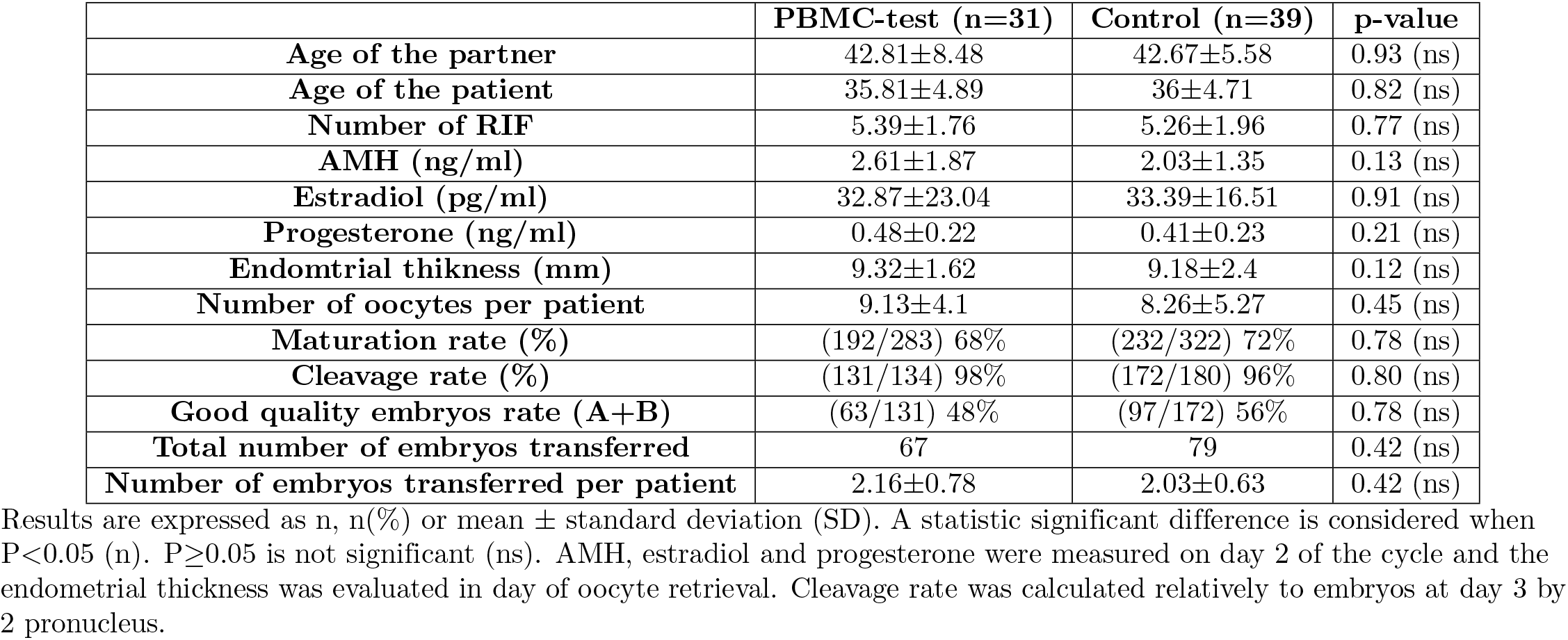
Comparison of the patient’s characteristics.

|  | PBMC-test (n=31) | Control (n=39) | p-value |
| --- | --- | --- | --- |
| Age of the partner | 42.81±8.48 | 42.67±5.58 | 0.93 (ns) |
| Age of the patient | 35.81±4.89 | 36±4.71 | 0.82 (ns) |
| Number of RIF | 5.39±1.76 | 5.26±1.96 | 0.77 (ns) |
| AMH (ng/ml) | 2.61±1.87 | 2.03±1.35 | 0.13 (ns) |
| Estradiol (pg/ml) | 32.87±23.04 | 33.39±16.51 | 0.91 (ns) |
| Progesterone (ng/ml) | 0.48±0.22 | 0.41±0.23 | 0.21 (ns) |
| Endometrial thickness (mm) | 9.32±1.62 | 9.18±2.4 | 0.12 (ns) |
| Number of oocytes per patient | 9.13±4.1 | 8.26±5.27 | 0.45 (ns) |
| Maturation rate (%) | (192/283) 68% | (232/322) 72% | 0.78 (ns) |
| Cleavage rate (%) | (131/134) 98% | (172/180) 96% | 0.80 (ns) |
| Good quality embryos rate (A+B) | (63/131) 48% | (97/172) 56% | 0.78 (ns) |
| Total number of embryos transferred | 67 | 79 | 0.42 (ns) |
| Number of embryos transferred per patient | 2.16±0.78 | 2.03±0.63 | 0.42 (ns) |

Results are expressed as n, n(%) or mean ± standard deviation (SD). A statistic significant difference is considered when $P < 0.05$ (n). $P \geq 0.05$ is not significant (ns). AMH, estradiol and progesterone were measured on day 2 of the cycle and the endometrial thickness was evaluated in day of oocyte retrieval. Cleavage rate was calculated relatively to embryos at day 3 by 2 pronucleus.

### IVF procedures

Embryos produced by ICSI [22] were cultured up to day 3. Adequate embryo quality (good quality embryos; A+B) was defined based on the presence of uniformly sized and shaped blastomeres and fragmentation lower or equal to 10%. One or two good quality embryos were transferred in utero using a Frydman catheter (CCD Laboratories, Paris, France). The implantation success (observation of the embryo sac) was assessed by ultrasound imaging and calculated relative to the number of transferred embryos. Clinical pregnancy was confirmed by ultrasound imaging 6-8 weeks after embryo transfer and calculated relative to the number of transferred cycles. The miscarriage ratio was calculated relative to the number of clinical pregnancies after the first trimester. Each couple went through a single ICSI cycle during this study.

### PBMC immunotherapy and LPL assay

After antagonist ovarian stimulation protocol, a blood sample is scheduled on the day of ovulation induction to isolate PBMCs using a separation protocol based on Ficoll. PBMCs are well prepared after a culture for 72 h and then transferred to the patient in utero two days before embryo transfer as it was elucidated by Madkour et al. [22]. After embryo transfer, patients receive oral Utrogestan (200 mg×2/day) for luteal support.

In the course of our study, the included patients underwent the LPL analysis at day 15 after embryo transfer to reflect the P4 synthesis by CL after implantation using the serum for the first pregnancy test for *β*-hCG assay. Indeed, the LPL analysis was assessed using the immunological technique of electro-chemiluminescence (ECLIA, Roche, Mannheim, Germany) at LABOMAC center.

### Statistical analysis

Data are presented as the mean ± standard deviation (SD) or percentage of the total. Data were analyzed with the Student’s t-test for comparison of mean values or with the chi squared test for comparison of percentages, and r-correlations using Statistical Package, version 6.0 (Statistica); p< 0.05 shows significant differences. Then, the mean values of each parameter’s results were evaluated to calculate the study power with the post-hoc test using the G*Power software (version 3.0.10).

## Results

Clinical outcomes were doubly improved after immunotherapy including implantation and clinical pregnancy in Control versus PBMC-test for patients with at least 3 RIF (10% and 21% vs 24% and 45%) while the effect of PBMC on miscarriage rate was non-significant (75% vs 21%; p=0.06). In the other hand, this strategy shows an increase over double in LPL (9ng/ml for PBMC-test vs 4ng/ml for Control) showing significant correlation with clinical pregnancy rate for PBMC treated patients (Table 2). Moreover, the LPL was not influenced by RIF number with non significant r correlation (r=−0.36).

**Table 2.** Clinical outcomes and luteal progesterone level after PBMC immunotherapy for patients with at least 3 RIF.

| Clinical outcomes | PBMC-test (n=31) | Control (n=39) | p-value | Power 1-β |
| --- | --- | --- | --- | --- |
| Implantation rate (%) | (16/67) **24%** | (8/79) **10%** | **0.02 (s)** | **89%** |
| Clinical pregnancy rate (%) | **(14/31) 45%** | **(8/39) 21%** | **0.03 (s)** | **71%** |
| Miscarriage rate (%) | (3/14) 21% | (6/8) 75% | 0.06 (ns) | **99%** |
| LPL (ng/ml) | **9.32±5.70** | **3.80±3.65** | **0.00001 (s)** | **99%** |
| r-correlation (LPL to clinical pregnancy) | **0.28 (p=0.01)** | 0.16 (p=0.32) | – | – |

Results are expressed as n, n(%) or mean ± standard deviation (SD). A statistic significant difference is considered when $P < 0.05$ (n). $P \geq 0.05$ is not significant (ns). Power 1- $\beta$ ( $\beta$ is error type II) is calculated basing on difference of mean values between two groups (PBMC-test vs Control) with $\alpha = 0.05$ ( $\alpha$ is error type I). Power value is considered highly important when it is above 80%. r-correlation was calculated relatively to LPL depending on clinical pregnancy rate in each group of PBMC-test and Control, and it is considered significant when $P < 0.05$ and non significant when $p \geq 0.05$ . The implantation rate is expressed as the ratio between the number of embryonic sacs and the total number of transferred embryos; the miscarriage rate is expressed relative to the number of clinical pregnancies. LPL (Luteal progesterone level) was measured at day 15 after embryo transfer.

## Discussion

Despite there is not clear differential clinical diagnostic to differ between RIF and RPL, over 75% of pregnancy failures are due to implantation failures [33]. Whatever controversies regarding RIF and RPL clinical definition, hypothetically in our previous study [22] it was suggested that RIF is due to pro-inflammatory (Th1) deficiency while RPL is due to Th1 persistence inhibiting the anti-inflammatory (Th2) release. Therefore, in this current study following the results of our previous work [22] PBMC immunotherapy was efficient for RIF patients since their second implantation failure to double their chance to conceive. Nevertheless, we are not the only team who are prescribing this kind of treatment to patients with RIF or generally with IVF failures. Yoshioka et al. were the pioneers in this approach application while the research teams’ followers could to involve some technical modifications in PBMC preparation protocol [23]. Some could to prove the efficiency of hCG supplementation on PBMC culture for 72h [22, 24] or trying to minimize latter to 24h [26, 27, 34] while others were more focused on the efficiency of CRH [25]. All these technical adaptations were occurred in order to enhance at maximum the function of PBMC and their cytokines secretions to activate thereafter the maternal immune system into endometrium after an intrauterine administration and be ready for embryo implantation. Indeed, as expected, implantation rate after PBMC immunotherapy was over double compared to control (24% vs 10%; table 2) and the result was similar to other studies with interval 21-25% for treated patients [22–24, 26, 27].

Nevertheless, it was commonly accepted that with insufficient P4 production causing miscarriages could be solved simply by an exogenous P4 administration in order to regulate the inflammatory mediators of pregnancy and even for patients undergoing IVF process in fresh or frozen cycles to improve clinical outcomes prior embryo transfer [29, 35, 36]. Indeed, P4 presents an anti-inflammatory action enhancing Th2 cytokines production to maintain pregnancy [37, 38].

However, our doubled clinical outcomes including implantation and clinical pregnancy rates could have more evident explanation especially when LPL showed high increase in PBMC-test compared to control (9ng/ml vs 4ng/ml) with positive correlation relatively to clinical pregnancy just for treated RIF patients (r=0.28)(Table 2). However, 4 ng/ml of LPL was not correlated to clinical pregnancy rates for RIF patients in control group (r=0.16). This observation allowed us to conclude that certainly even with P4 importance to maintain clinical pregnancy, this latter could be occurred. Moreover, it explained why P4 supplementation treatments kept into question their efficiency for RIF, RPL and RM patients despite the maternal immune system is dysregulated led toward Th1 or Th2 [21, 29, 39, 40]. In the opposite side, when this latter is probably trying to turn back its balance via PBMC immunotherapy which acted not only into endometrium but also in CL to produce more the P4, LPL became correlated to clinical pregnancy. This hypothesized interactive reaction between LPL issued from CL and PBMC administrated into endometrium, is realized in order to assure the embryo implantation and pregnancy maintain homing Th1/Th2 balance(Fig 1).

**Fig 1.**
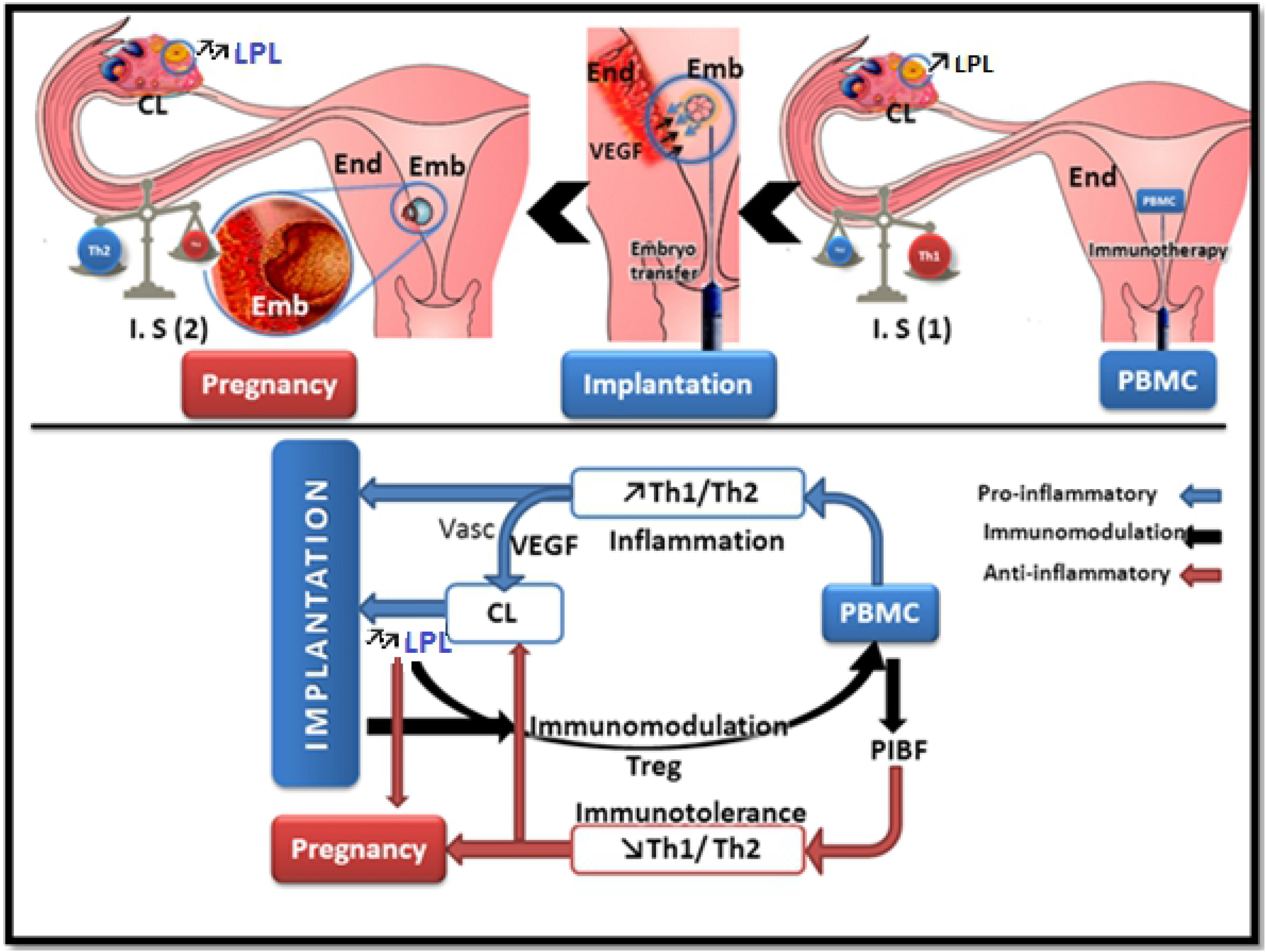
Hypothesis of interactive effect of PBMC immunotherapy and luteal progesterone level for implantation and pregnancy success. After intrauterine administration PBMC, the Th1 / Th2 balance towards Th1 tends to ensure ignition shift by secreting pro-inflammatory cytokines and several growth factors primarily VEGF inducing vascularization in one hand into endometrium to prepare for embryo invasion embryo and in the other hand into CL in order to increase luteal progesterone level (LPL). All these immune-endocrine factors are limited in closest communication circle with mutual interactive modulation to ensure the embryo implantation. Thereafter, an increased LPL can induce immunomodulation by promoting T cells differentiation into Treg and secreting PIBF as an immunosuppressor factor that promotes the Th1/Th2 balance to Th2 anti-inflammatory system ensuring immunotolerance of allograft “embryo”. Thus, Th2 cytokines secretion involved in CL maturation can eventually to increase more LPL required for pregnancy maintain. CL: Corpus Luteum; LPL: Luteal progesterone level; I. S (1): Immune System (Pro-inflammatory); I. S (2): Immune System (Anti-inflammatory); PBMC: Peripheral Blood Mononuclear Cells; End: Endometrium, Emb: embryo; Vasc: Vascularization; PIBF: Progesterone induced blocked factor; VEGF: Vascular Epidermal Growth Factor, Treg: Lymphocyte T regulator.

Furthermore, luteal P4 would regulate the uterine level synthesis of CSF-1, cytokine essential for the vascularization of the endometrium and to maintain pregnancy by increasing just before implantation to achieve a peak tripled to day 15 of pregnancy [41, 42]. Our results show a 63% increase in the synthesis of the luteal progesterone in pregnant patients treated with PBMC which joins perfectly the observed effect of PBMC on clinical pregnancy rate in our study. It seems that this effect is mediated by Th2 cells secreting IL4 and IL10 able to optimize the recruitment of leukocytes for VEGF secretion in CL [13]. The latter is better vascularized release P4 production [43].

Te PBMC effect could not be certainly over early pregnancy stage to balance maternal immune system and LPL while P4 will be placental and the ongoing pregnancy until delivery will be more influenced by fetus and genetic reproductive function. May be for this reason, PBMC immunotherapy is less effective to avoid miscarriages as showed in our study with non-significant difference in miscarriage rate (21% for PBMC-test and 75% for control; Table 2) and confirmed by others [22, 26].

The paramount function of PBMC is to provide trophic support for endometrium to be decidualized and for formation of dense vascular network in CL to produce more P4 that is essential for pregnancy maintain. However, perturbations of immune-endothelial cell crosstalk within the ovary during the peri-conceptional period are likely to be pivotal in luteal insufficiency in women. This issue could provide more therapeutic trends to enhance luteal function through the targeting of immune system.

## Conclusion

This immunotherapeutic strategy based on PBMC intrauterine administration suggests that embryo implantation is controlled by maternal immune cells in utero and this treatment is showed its efficiency for RIF patients doubling their clinical outcomes with significant increase of LPL. This issue demonstrated that immunotherapy had positive effect on luteal P4 synthesis during implantation which acted dually on homing maternal immune system into endometrium in order to maintain pregnancy. This non-invasive and much less expensive treatment than the multiplication of IVF attempts could be proposed as part of ART to patients since their second implantation failure or even for patients with RPL or RM who are directly redirected to be treated with P4 supplementation or other anti-inflammatory treatments. Nevertheless, this issue needs an eventual researches and clinical investigations.

## Acknowledgments

The authors acknowledge the help and expertise of the research team of Anfa Fertility Center at Embryology laboratory and LABOMAC clinical laboratory.

